# Early and lifelong effects of APOE4 on neuronal gene expression networks relevant to Alzheimer’s disease

**DOI:** 10.1101/2022.06.16.496371

**Authors:** Brian P. Grone, Kelly A. Zalocusky, Yanxia Hao, Seo Yeon Yoon, Patrick Arriola, Yadong Huang

**Author notes:** Correspondence: Yadong Huang or Brian Grone.

## Abstract

Apolipoprotein E4 (*APOE4*) genotype and aging are critical risk factors for Alzheimer’s disease (AD). Aged APOE4 knock-in (APOE4-KI) mice have phenotypes reflecting features of AD. We conducted a large-scale single nucleus RNA-sequencing study to identify cell-type-specific effects of APOE4 on hippocampal gene expression during aging. APOE4-KI mice showed prominent alterations, relative to APOE3-KI mice, in neuronal transcriptome related to synaptic function, calcium signaling, and MAPK/Rap1/Pld signal transduction, starting by 5 months and persisting during aging. Mice with the *APOE4* gene removed specifically from neurons failed to show most of these neuronal transcriptomic changes, suggesting a specific effect of neuron-derived APOE4 on the transcriptome. APOE4 affects similar cellular pathways in induced pluripotent stem cell-derived human neurons transplanted into APOE4-KI mouse hippocampus and in cortical neurons from aged human brains. Thus, neuronal APOE4 has early and persistent effects on neuronal transcriptomes, suggesting the requirement of early interventions for successfully treating *APOE4*-related AD.

## Introduction

Alzheimer’s disease (AD) is a progressive neurodegenerative disorder that leads to memory loss and severe cognitive impairment. Currently, AD is a prevalent and growing cause of death and disability, with no treatments available that can alter the course of the disease^1,2^. As AD progresses, brain structure and function deteriorate, with synapse loss and neuron death in hippocampus, cortex, and other brain regions^3^. The ε4 allele of apolipoprotein E (*APOE4*) is the major genetic risk factor for late onset AD. Carrying the *APOE4* allele increases the risk of AD in a gene dose-dependent manner compared to the *APOE3* allele^4-6^. The impact of APOE4’s large effect size is magnified by its prevalence in the global human population. Among AD patients, *APOE4* heterozygous carriers account for ∼40–60% of AD patients, and *APOE4* homozygous carriers account for ∼5–14%, with rates varying by country and region^7^. The other major risk factor for AD is advanced age, with the risk of AD approximately doubling every 5 years after age 65^8^. However, it is not clear how APOE4 and aging effects are related, and how they interact to elicit disease phenotypes. Understanding the specific pathways that mediate APOE4 contributions to AD pathogenesis could allow targeted development of interventions to treat *APOE4*-related AD or reduce AD risk for APOE4 carriers^9,10^. Bulk RNA-sequencing (RNA-seq) studies in mouse models have identified some genes and pathways that are altered by APOE genotype and age^11^. Furthermore, studies of iPSC models in culture have demonstrated APOE genotype effects in cell types including neurons, astrocytes, microglia, and pericytes^12,13^. However, it is still not clear which cell types are specifically affected by APOE genotype in an *in vivo* condition with all cell types together, and at what ages the effects begin to occur.

Early APOE4 effects on brain and behavior have been reported in humans and mice before the onset of severe disease symptoms^14^. APOE4 knock-in (APOE4-KI) mice have delayed development of neurons in embryonic and postnatal stages^15^, lower dendritic spine density beginning as early as 4 weeks^16^, and impaired long-term potentiation (LTP) at 2–4 months^17^. APOE4-KI mice also show impaired learning and memory as early as 3 to 4 months of age in some behavioral tests^18-20^. In humans, young adult *APOE4* carriers may have hippocampal hyperactivity^21^ and white matter deficits relative to non-carriers^22^, and APOE4 is associated with memory declines before age 60, even in carriers without MCI or AD^23,24^. Despite extensive evidence for a variety of early effects of APOE4, the timing and progression of its transcriptome-wide effects in specific brain cell types remains understudied.

The molecular and cellular mechanisms that lead to APOE4-associated deficits, and eventually to neurodegeneration and disease, are not well understood. However, important aspects of the location and timing of APOE4 effects have been identified in humans and in mouse models. Although astrocytes are a predominant cell type expressing APOE in the brain, some neurons also express APOE^25-28^. Stresses during aging may lead to APOE expression in neurons^29,30^, resulting in tau-dependent neurotoxicity as well as synaptic and neuronal degeneration^25-28^. Recently, it was shown that APOE expression level in neurons is correlated with immune-response gene expression, particularly MHC-I, which leads to tau-related neuropathology and neuronal death^31^. Inhibitory neurons show deficits in APOE4-KI mice, where their dendritic arborization and synaptic density are diminished^32,33^. GAD67-positive neurons are progressively lost after 3 months in female APOE4-knockin mice, compared to APOE3-KI mice^34^. Excitatory synaptic activity is also impaired in APOE4 mice^35^ and progressively worsens^36^. Despite decades of research on APOE4, however, previous studies have not been able to systematically identify which cells and genes are involved in its effects and at what stage of disease or development they are affected.

To understand how APOE4 affects the various types of hippocampal cells throughout adulthood, we analyzed single-nucleus RNA-sequencing (snRNA-seq) data from the hippocampus of APOE4-KI and APOE3-KI mice aged 5, 10, 15, and 20 months. We characterized gene expression patterns affected by APOE4, at each age, in a cell-type-specific manner and identified prominent pathways affected by APOE4 throughout the lifespan. Our data show that APOE4 disrupts key neuronal gene networks that regulate synaptic activity, calcium signaling, and MAPK/Rap1/Pld pathways. Furthermore, evidence from mice with the *APOE* gene specifically deleted in neurons suggests that APOE4 effects on neuronal transcriptomes may depend on APOE expression within neurons themselves, even though APOE expression is found at higher levels in astrocytes and other cell types. Importantly, we found that effects of APOE4 on neuronal transcriptome are apparent early in adulthood and persist during aging. In contrast, although some non-neuronal cell types, particularly astrocytes and oligodendrocytes, also show significant effects of APOE4 on their transcriptomes, they occurred at older ages. Finally, similar effects of APOE4 on neuronal transcriptome are found in induced pluripotent stem cell (iPSC)-derived human neurons transplanted into APOE4-KI mouse hippocampus and in cortical neurons from aged human brains, supporting potential translation of these experimental observations from mice to humans.

## Results

### The transcriptomic effects of APOE4 begin early in neurons and late in non-neuronal cells in mouse hippocampus

To examine differential transcriptomic effects of human APOE4 and APOE3, we used a snRNA-seq dataset (GEO:GSE167497) that has been previously analyzed in a separate study in our lab^31^. For this dataset, hippocampi were collected from female APOE3-KI and APOE4-KI mice at 5, 10, 15, and 20 months of age (n = 4 per genotype and age) for snRNA-seq (Fig. 1a). Female mice were used owing to their susceptibility to APOE4-induced neuronal and behavioral deficits during aging^34^. For the present study, we used the previously reported^31^ filtering, clustering, and cell type classifications (Fig. 1b). This data set consists of 123,489 nuclei, including 16 distinct neuronal clusters (105,644 nuclei) and 11 non-neuronal clusters (17,845 nuclei). Each age and genotype shows distribution of nuclei across excitatory and inhibitory neuron clusters, astrocytes, oligodendrocytes, microglia, endothelial cells, and other cell types (Fig. 1c). We reanalyzed this original dataset without imputation of RNA expression level.

**Fig. 1.**
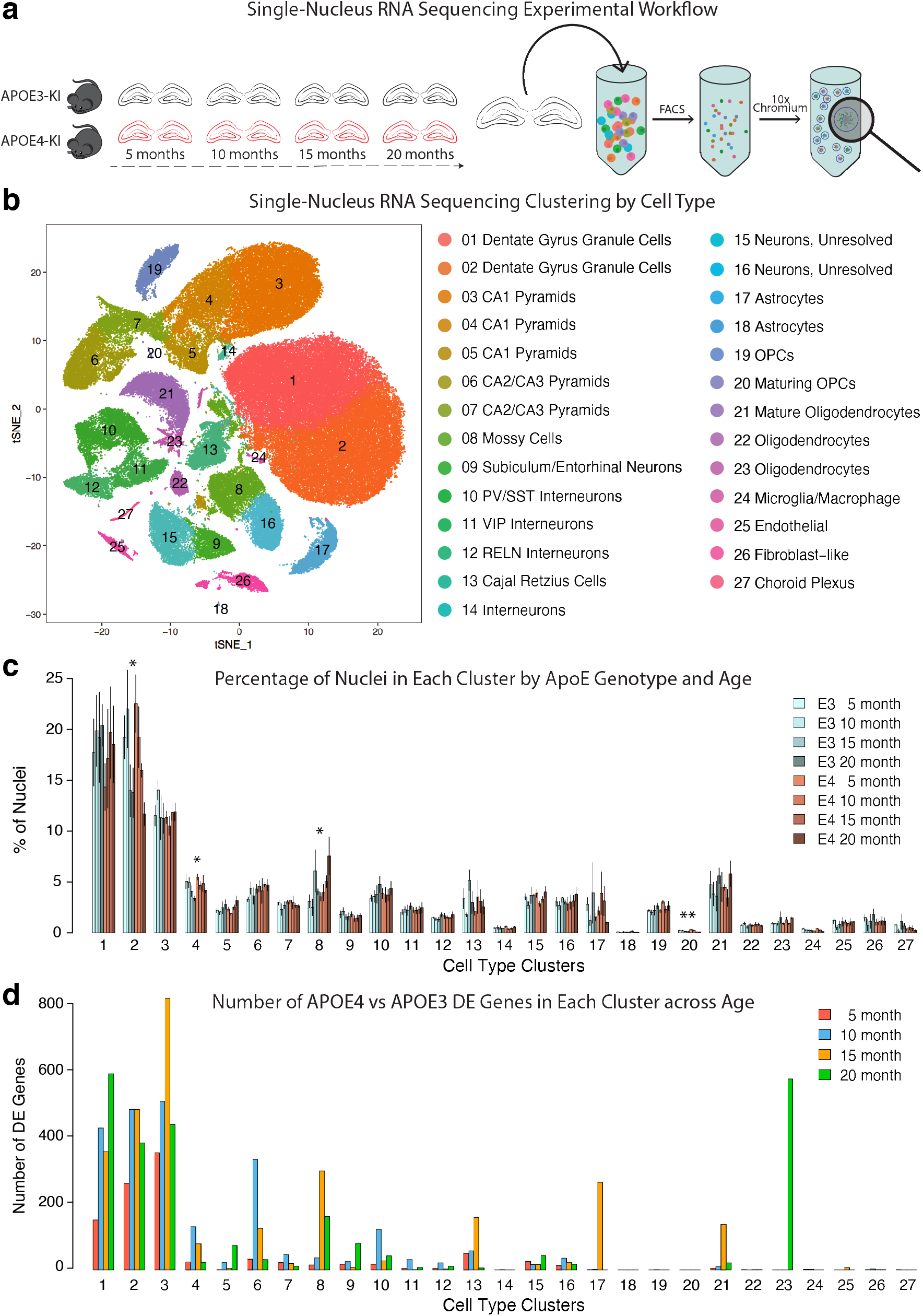
The transcriptomic effects of APOE4 begin early in neurons and late in non-neuronal cells in mouse hippocampus. **a**, Hippocampi were dissected from female APOE3-KI and APOE4-KI mice at 5, 10, 15, and 20 months of age (n = 4 per genotype per age). The hippocampi were dissociated, and nuclei were labeled with DAPI and isolated using flow cytometry before processing with the 10x Chromium v2 system (GEO:GSE167497). **b**, Clustering using the Seurat package revealed 27 distinct cell clusters. Marker gene analysis led to the identification of 16 neuronal clusters (Clusters 1–16) and 11 non-neuronal clusters (Clusters 17–27). **c**, Cell clusters showing the mean value of cell numbers from four individual samples in each group. Lines above the bars show size of standard errors (SE). Asterisks indicate significant effects of age on cell numbers by two-way ANOVA test (*<0.05, **<0.005; for clusters 2 (Pr(>F)= 0.00602), 4 (Pr(>F)= 0.0471), 8 (Pr(>F)= 0.0481), and 20 (Pr(>F)= 0.000467). **d**, The number of DE genes (on the y-axis) detected between APOE4-KI and APOE3-KI samples is shown for each cluster at each age (on the x-axis).

Two-way ANOVA was used to test potential effects of age, genotype, and age:genotype interaction on cell numbers. Three cell clusters showed a statistically significant decrease in cell numbers with age, including clusters 2 (dentate gyrus granule cell), 4 (CA1 pyramidal neuron), and 20 (maturing OPC) (Fig. 1c). Cluster 8 (mossy cell) had a statistically significant effect of age, with increasing numbers at older ages (Fig. 1c). The two-way ANOVA tests detected no significant effects of *APOE* genotype on cell numbers and no significant interaction effects between age and *APOE* genotype.

For each cluster of cells, differentially expressed (DE) genes between APOE4-KI and APOE3-KI mice were assessed at each age using zinbwave and edgeR packages in R. DE genes were detected in most neuron types (clusters 1–16) from age 5 months onwards, and the DE gene numbers increased at later ages (Fig. 1d and Supplementary Table 1). At each age, *APOE4* genotype shows more downregulated genes than upregulated genes (Supplementary Table 1). The numbers of DE genes varied across neuronal clusters (Fig. 1d) and were generally highest in the dentate gyrus granule cells (Clusters 1 and 2) and hippocampal CA1 pyramidal neurons. In contrast, DE genes were detected in non-neuronal cell clusters, particularly astrocytes and oligodendrocytes, after 15 months of age (Fig. 1d). Thus, the transcriptomic effects of APOE4 begin early in neurons and late in non-neuronal cells in this data set. To test the potential confounding effect of cell number variations across clusters for the DE gene analyses, we down-sampled the cells to a maximum of 500 or 1000 per genotype in each cluster at each age and tested differential expression. The pattern of DE gene numbers persisted even when subsamples of cells were used (Extended Data Fig. 1), suggesting that the cell number variations do not confound the analyses of APOE4’s effects on transcriptomes.

### APOE4 induces consistent DE pathways in hippocampal neurons across ages

The DE genes detected at each age (5, 10, 15, and 20 months) for each cluster (Fig. 1d and Supplementary Table 1) were used as input to determine significantly enriched KEGG (Kyoto Encyclopedia of Genes and Genomes) pathways (Supplementary Table 2). All pathways that were in the top 10 most enriched KEGG pathways for any cluster from 5 to 15 months were shown as heatmaps (Fig. 2a–c). Similar pathways were also affected by APOE4 at 20 months (Extended Data Fig. 2). Increasing numbers of DE pathways were significantly affected as age increased, from 23 (5 months) to 67 (20 months) (Supplementary Table 2).

**Fig. 2.**
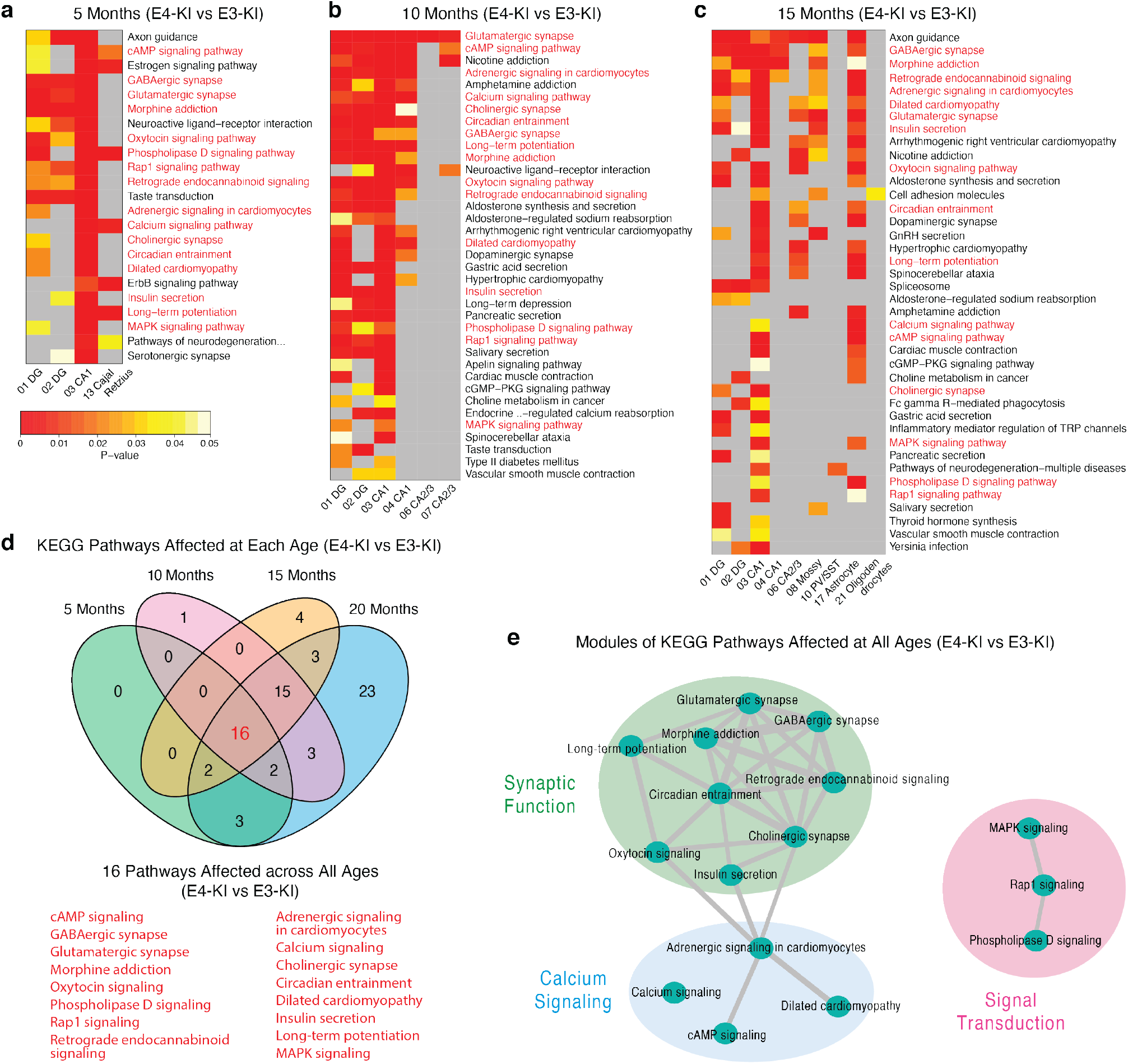
APOE4 induces consistent DE pathways in hippocampal neurons across ages. **a–c**, DE genes of APOE4-KI versus APOE3-KI mouse hippocampal cell types are significantly enriched in several KEGG pathways at 5 months (a), 10 months (b), and 15 months (c). Heatmap plots show cell clusters with significantly enriched DE pathways (on the x-axis) and the top enriched DE pathways (on the y-axis) sorted from most commonly detected (top) to least (bottom). P-value for statistical enrichment is shown by color, with red indicating smaller P-value, gray indicating P-value > 0.05, and scale indicated in the legend. Pathway names highlighted in red are found at all ages. **d**, A Venn diagram shows the number of shared DE pathways between each combination of ages (5, 10, 15, and 20 months). The 16 pathways enriched at all four ages are shown in the list. **e**, Network graph of the 16 DE pathways enriched at all four ages. The percent overlap of genes between pathways is represented by line thickness.

We observed that the most enriched DE pathways were found in neuron clusters (Clusters 1–16), starting as early as 5 months (Fig. 2a). In particular, excitatory neuron clusters showed a large number of DE pathways, with Cluster 3 (CA1 pyramidal Neurons) showing many highly enriched pathways at each age (Fig. 2a–c and Extended Data Fig. 2). Astrocytes (Cluster 17) showed enrichment in several similar DE pathways at 15 months. To examine how similarly pathways were affected by APOE4 across ages, we made a Venn diagram showing the number of DE pathways shared between each combination of ages (Fig. 2d). As in the heatmap plots, only those pathways in the top 10 significantly enriched DE pathways for any cell cluster were included in the Venn diagram. This analysis revealed 16 shared DE pathways across all ages (Fig. 2d and pathways highlighted in red in Fig. 2a–c and Extended Data Fig. 2).

To examine relationships between these 16 shared DE pathways affected by APOE4, we used genes shared across the pathways to construct a graph of the similarity of the 16 DE pathways. We then calculated percent overlap of genes between clusters and plotted the resulting network of pathways. This analysis of DE pathway similarity allowed us to identify 3 modules of highly interconnected pathways affected by APOE4: Synaptic Function (green), which was the module with the most included pathways, Calcium Signaling (blue), and MAPK/Rap1/Pld Signal Transduction (pink) (Fig. 2e).

### APOE4-induced DE genes in the shared pathways in hippocampal neurons across age

We focused our further analyses on excitatory neurons, which showed the largest numbers of DE genes and pathways starting as early as 5 months (Fig. 1d, Fig. 2a–c, and Supplementary Table 1 and 2), and examined the relationships between these pathways in more detail. The DE genes in the 16 shared DE pathways were analyzed for the larger excitatory neuron clusters 1–3 at each age. Heatmaps show the relationships of DE genes and pathways for Cluster 3 (CA1 pyramidal neuron) (Fig 3a–c) and Clusters 1 and 2 (dentate gyrus granule cells) (Extended Data Fig. 3 and 4).

**Fig. 3.**
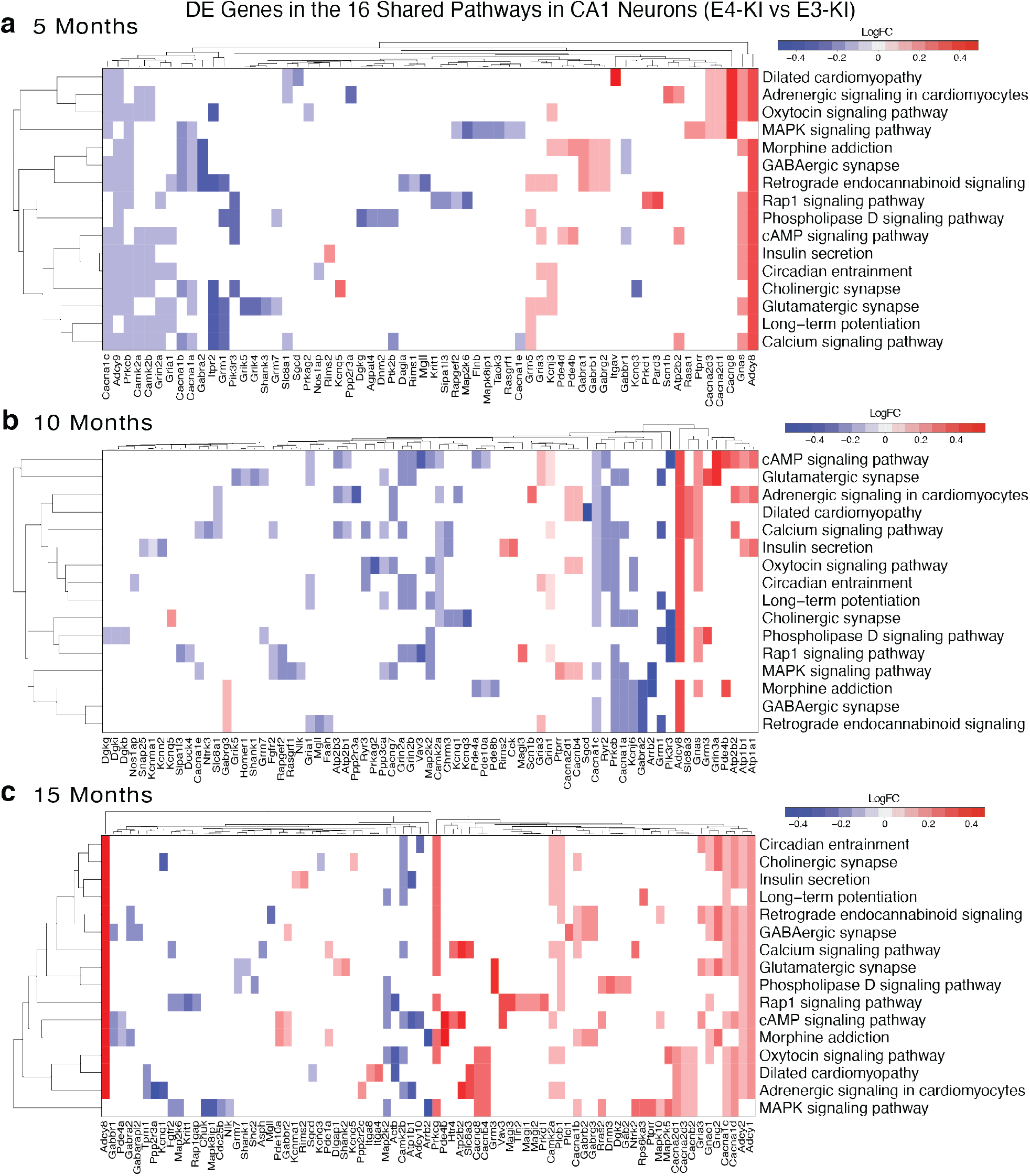
APOE4-induced DE genes in the 16 shared DE pathways in hippocampal CA1 pyramidal neurons across age. **a–c**, The DE genes in 16 shared pathways affected by APOE4 were plotted as heatmaps at 5 months (a), 10 months (b), and 15 months (c). For this analysis, we used the enrichKEGG() function, and included KEGG pathways with minGSSize = 10, pvalueCutoff = 0.05, Count min = 4. Genes were included if FDR < 0.05 and logFC > 0.1. The APOE4 effect, relative to APOE3, is plotted as intensity of blue (downregulation) or red (upregulation), scale indicated in the legend. DE genes are clustered on the x-axis, and pathways are clustered on the y-axis, by similarity of expression effects.

The results showed that APOE4-KI mice at each age had up-and down-regulated genes across all the 16 shared DE pathways in CA1 pyramidal neurons and dentate gyrus granule cells. Furthermore, some DE genes were found in several of the 16 shared DE pathways (Fig. 3a–c and Extended Data Fig. 3 and 4), suggesting that they could be responsible for multiple aspects of the APOE4 effects in neurons. These shared DE genes largely belong to the categories of ion channels, GABA receptors, phosphatases, protein kinases, or adenylate cyclases (Fig. 3a–c and Extended Data Figs. 3 and 4). Strikingly, many of these DE genes shared across the 16 DE pathways were also shared across age, starting as early as 5 months (Fig. 3a–c and Extended Data Fig. 3 and 4), suggesting their potential early and persistent roles in APOE4’s contributions to AD pathogenesis.

### Neuron-specific deletion of the *APOE4* gene identifies neuronal APOE4-induced DE pathways and modules across age

We then examined the effects of neuron-derived APOE isoforms on hippocampal transcriptomes using lines of APOE-KI mice carrying the human *APOE3* or *APOE4* gene flanked by loxP sites^37^. To remove the human *APOE* gene from neurons, these mice were crossed with mice expressing the Cre recombinase driven by the neuron-specific Synapsin 1 promoter (Syn1-Cre). Hippocampal samples were collected from APOE4-fKI/Syn1-Cre and APOE3-fKI/Syn1-Cre mice at 5, 10, and 15 months. The snRNA-seq data from 5-month and 10-month APOE-fKI/Syn1-CRE mice were newly acquired for this study. Samples were collected and analyzed using the same Seurat and downstream expression pipeline that we used for the APOE-KI dataset (Extended Data Fig. 5). The snRNA-seq data were filtered, normalized, and clustered, and cell types were identified using positively enriched marker genes, as we reported previously^31^. The snRNA-seq data and clustering of 15-month-old APOE-fKI/Syn1-CRE mice were taken from our previously reported dataset (GEO:GSE167497)^31^.

For the 5-, 10-, and 15-month datasets, DE genes in each cell cluster of the hippocampus were detected by comparing the APOE4-fKI/Syn1-Cre mice with the APOE3-fKI/Syn1-Cre mice using zinbwave and edgeR packages in R (Supplementary Table 3). We reasoned that the genes which were differentially expressed despite the absence of APOE in neurons may be regulated by APOE produced in brain cell types other than neurons. Therefore, for further analyses, we chose to remove such genes from our DE gene list derived from the comparison between APOE4-KI and APOE3-KI mouse hippocampus. This left a set of DE genes that are differentially expressed between APOE4 and APOE3 genotypes only when the *APOE* gene was present in neurons (as well as other cell types), which we referred to as neuronal APOE4-induced DE genes (Supplementary Table 4).

KEGG pathway enrichment analysis with the neuronal APOE4-induced DE genes revealed that, similar to the APOE4-KI versus APOE3-KI data analysis, most neuronal APOE4-induced DE pathways were found in neuron clusters, particularly excitatory neuron clusters, across age, again starting as early as 5 months (Fig. 4a–c and Supplementary Table 5). To examine how similarly pathways were affected across ages, we again made a Venn diagram showing the number of neuronal APOE4-induced DE pathways shared between each combination of ages (Fig. 4d). Strikingly, the same 16 shared DE pathways as those in APOE4-KI versus APOE3-KI comparison were identified (Fig. 4d), indicating that these 16 DE pathways are affected predominantly by neuronal APOE4.

**Fig. 4.**
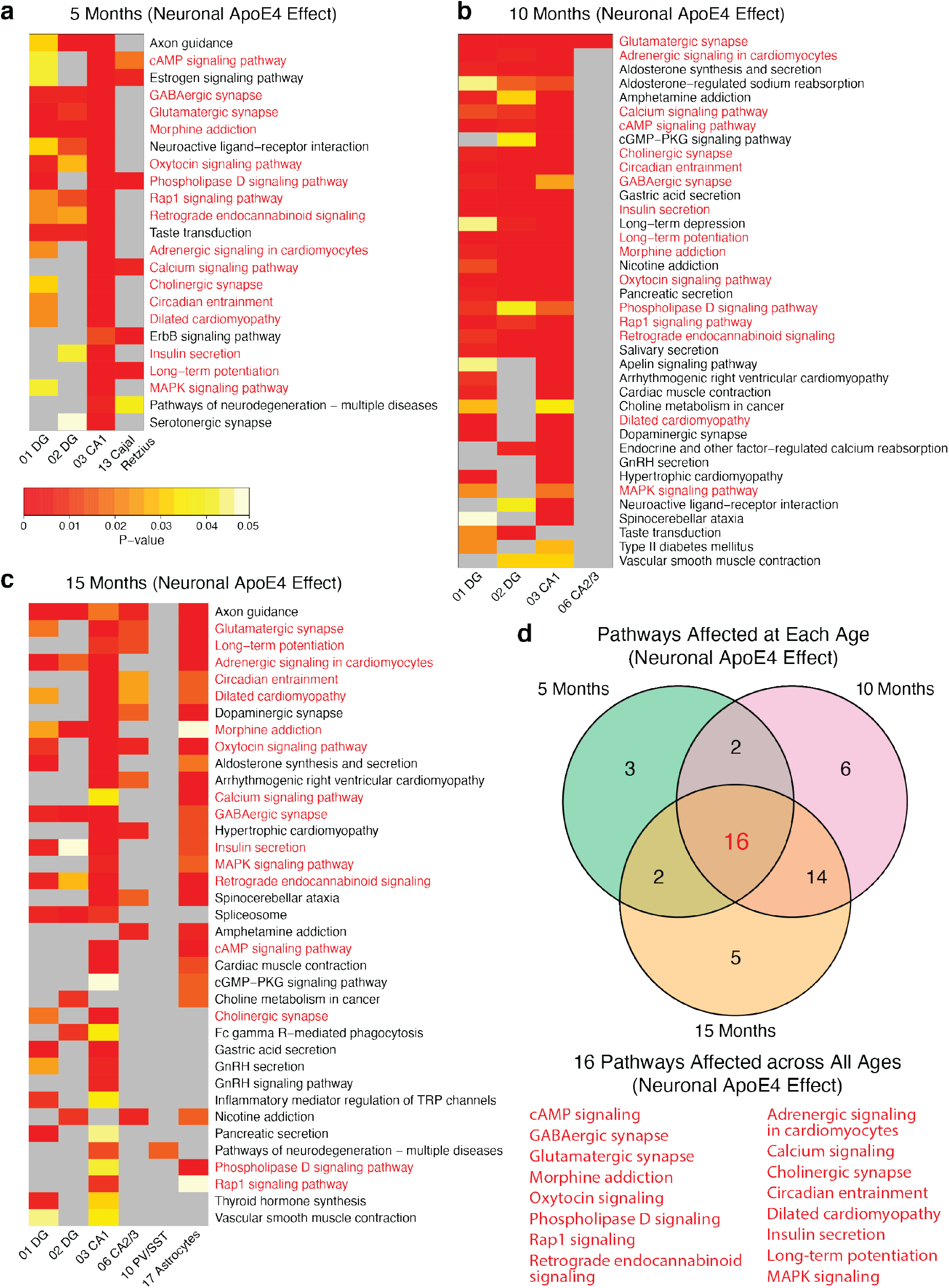
Neuron-specific deletion of the *APOE4* gene identifies neuronal APOE4-induced DE pathways and modules across age. **a–c**, Genes that are differentially expressed even in the absence of *APOE* gene in neurons were removed from the DE gene list derived from the comparison between APOE4-KI and APOE3-KI mouse hippocampus. The remaining gene list includes neuronal APOE4-induced DE genes that were significantly enriched in several KEGG pathways at 5 months (a), 10 months (b), and 15 months (c). Heatmap plots show cell clusters with significantly enriched neuronal APOE4-induced DE pathway (on the x-axis) and the top DE pathways (on the y-axis) sorted from most commonly detected (top) to least (bottom). P-value for statistical enrichment is shown by color, with red indicating smaller P-value, gray indicating P-value > 0.05, and scale indicated in the legend. Pathways with names highlighted in red were found at all ages. **d**, A Venn diagram shows the number of shared neuronal APOE4-induced DE pathways between each combination of ages (5, 10, 15 months). The 16 pathways enriched at all three ages are shown in the list.

### Neuronal APOE4-induced DE gene networks in major types of hippocampal neurons across age

Using the neuronal APOE4-induced DE gene list, we examined the relationships of these genes in the 16 shared DE pathways across age. The gene expression network analysis showed that the neuronal APOE4-induced DE genes included both up-and down-regulated genes across all the 16 shared DE pathways in CA1 pyramidal neurons (Fig. 5a–c) and dentate gyrus granule cells (Extended Data Fig. 5 and 6) across age. Furthermore, some DE genes were found in several of the 16 shared DE pathways (Fig. 5a–c and Extended Data Fig. 6 and 7), suggesting that they could be responsible for multiple aspects of neuronal APOE4 effects. Again, these shared DE genes largely belong to the categories of ion channels, GABA receptors, phosphatases, protein kinases, or adenylate cyclases (Fig. 5a–c and Extended Data Fig. 6 and 7). Again, many of these DE genes shared across the 16 DE pathways were also shared across age, starting as early as 5 months (Fig. 5a– c and Extended Data Fig. 6 and 7), suggesting their potential early roles in neuronal APOE4’s contributions to AD pathogenesis.

**Fig. 5.**
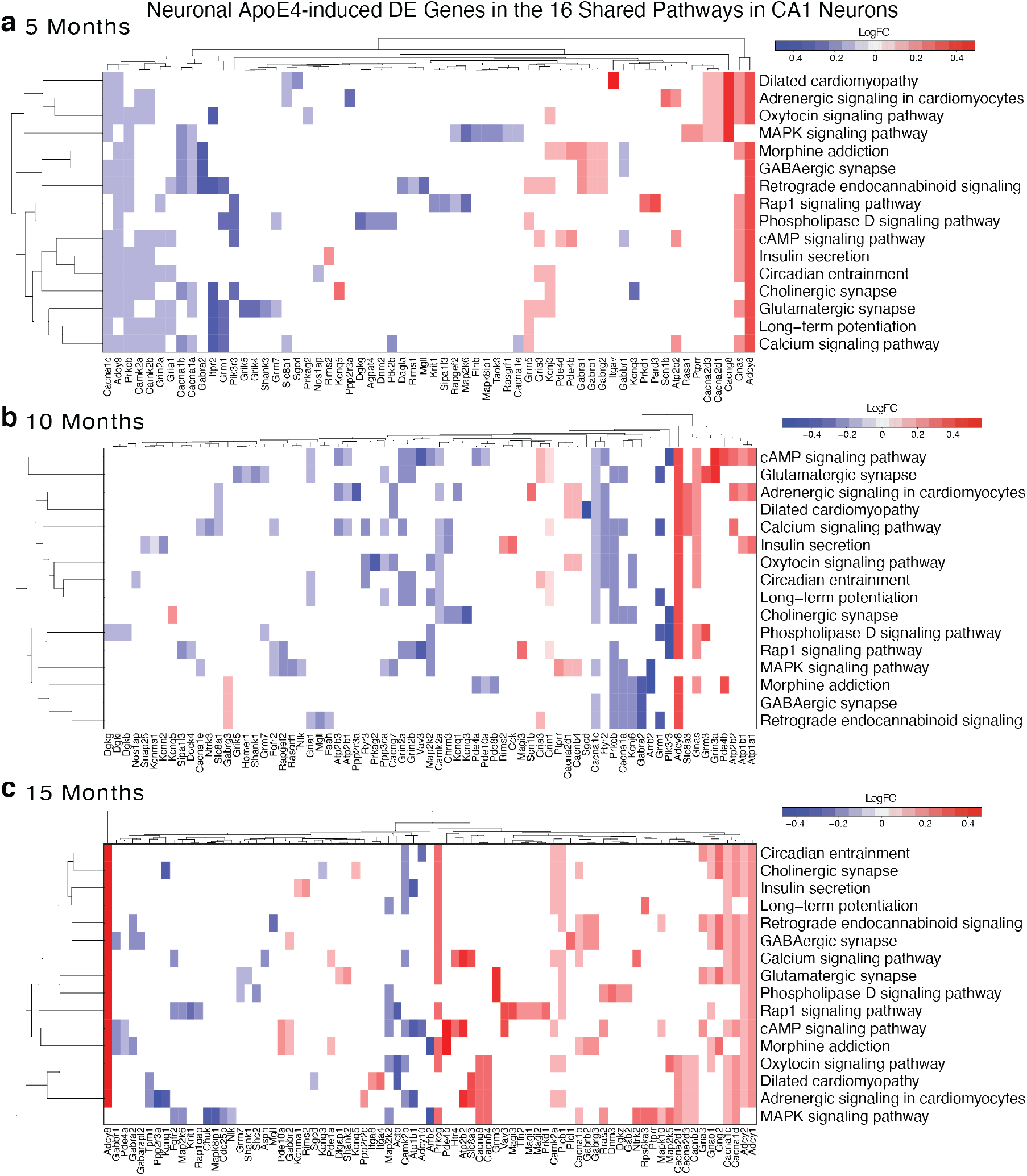
Neuronal APOE4-induced DE genes in hippocampal CA1 pyramidal neurons across age. **a–c**, The CA1 pyramidal neurons (cluster 3) revealed genes that were differentially expressed in APOE4-KI versus APOE3-KI mice, but not in APOE4-fKI/Syn1-Cre versus APOE3-fKI/Syn1-Cre mice. These neuronal APOE-affected DE genes that fell into 16 consistently affected pathways were plotted as heatmaps at 5 months (a), 10 months (b), and 15 months (c). For this analysis, we used the enrichKEGG() function, and included KEGG pathways with minGSSize = 10, pvalueCutoff = 0.05, Count min = 4. Genes included were DE genes between APOE4-KI and APOE3-KI mice, with genes removed if they were also DE genes between APOE4-fKI/Syn1-Cre and APOE3-fKI/Syn1-Cre mice. For each DE gene, the expression effect of neuronal APOE4 versus APOE3 was plotted as intensity of blue (downregulation) or red (upregulation).

### Neuronal APOE4-induced DE genes and pathways are shared by human APOE4 iPSC-derived neurons transplanted into APOE-KI mouse hippocampus

To test the relevance of the identified neuronal APOE4-induced DE gene and pathways to human neurons, we compared the mouse neuronal DE genes and pathways affected by APOE4 in APOE4-KI versus APOE3-KI mouse hippocampus with those obtained from human APOE4 and APOE3 iPSC-derived neurons transplanted into APOE-KI mouse hippocampus (referred to as an *in vivo* chimeric human neuron model)^38^. The raw snRNA-seq data (GEO:GSE152867) of human APOE4 and APOE3 iPSC-derived neurons transplanted into APOE-KI mouse hippocampus were the same as used in a previously published study from our lab^38^.

Seurat analysis and clustering of the human APOE4 and APOE3 iPSC-derived cells from the *in vivo* chimeric model identified four major cell types, including excitatory neuron, inhibitory neuron, astrocyte, and other (Fig. 6a). For our analysis, we used the APOE4 and APOE3 iPSC-derived cells exclusively from transplants into APOE4-KI mice. Analysis of the DE genes affected by APOE4 in human excitatory neurons in the APOE4-KI mouse background (Supplementary Table 6) identified the top 15 DE pathways (Fig. 6b and Supplementary Table 7). Nine out of the 15 DE pathways in human excitatory neurons overlapped with the 9 out of 16 DE pathways in mouse hippocampal excitatory neurons (Fig. 6c). Gene network analysis revealed the DE genes across the 9 shared DE pathways in human excitatory neurons in the APOE4-KI mouse background (Fig. 6d). Together, these data indicate consistency in transcriptomic effects of neuronal APOE4 between mouse neurons and human iPSC-derived neurons.

**Fig. 6.**
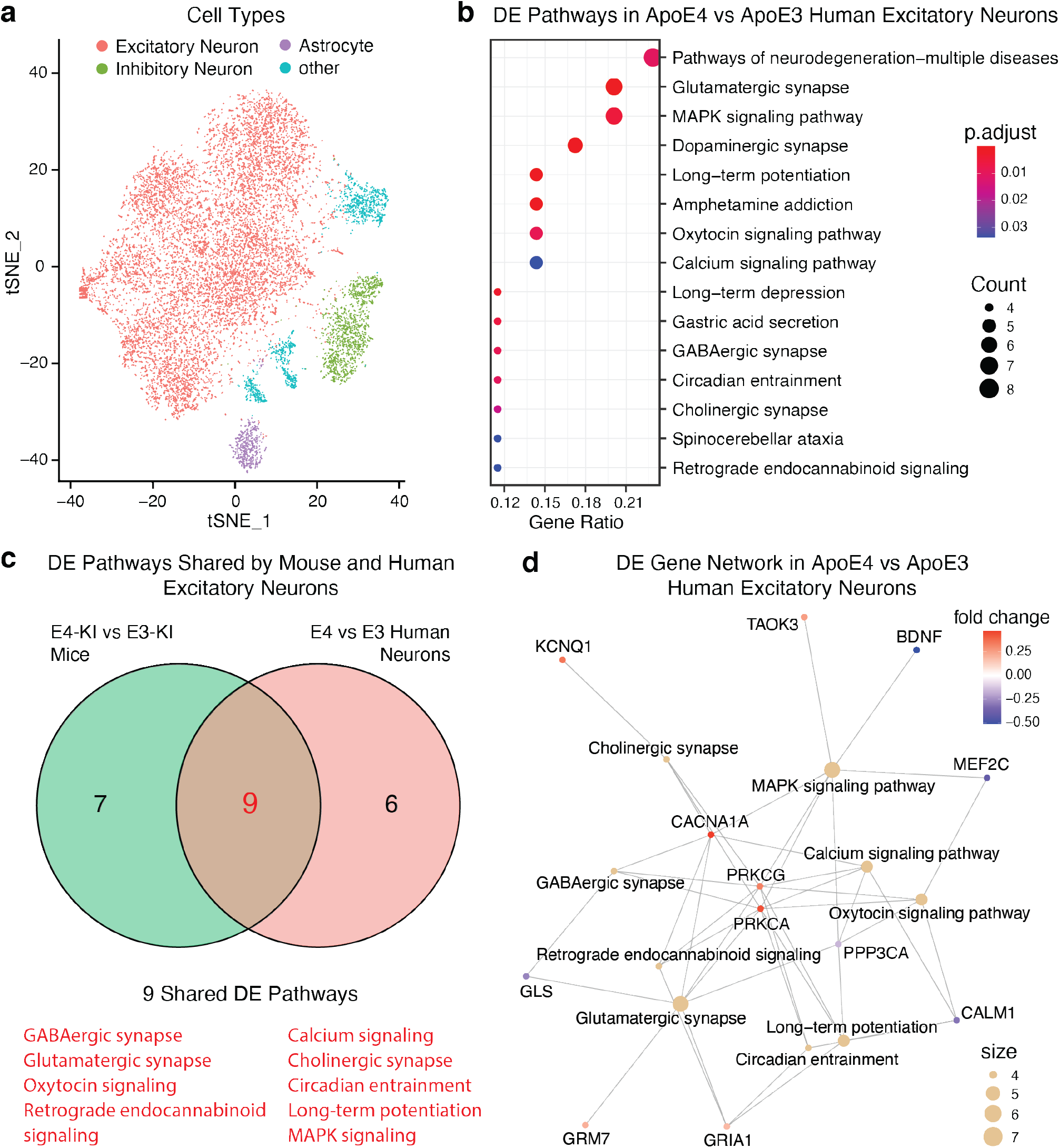
Neuronal APOE4-induced DE genes and pathways are shared by human APOE4 iPSC-derived neurons transplanted into APOE4-KI mouse hippocampus. **a**, tSNE plot shows major cell clusters, including excitatory neurons, inhibitory neurons, astrocytes, and other cells, derived from human APOE4 and APOE3 iPSCs, transplanted into APOE4-KI mice (GEO:GSE152867). **b**, Excitatory neurons were analyzed for DE genes between human APOE4 and APOE3 iPSC-derived excitatory neurons transplanted into APOE4-KI mice. The DE genes were further tested for KEGG pathway enrichment, and the top 15 DE pathways are shown. ‘Count’ shows number of DE genes identified in a pathway, corresponding to size of the dot; ‘p.adjust’ shows adjusted P-value for the statistical enrichment of genes in a pathway, red color corresponds to lower P-value. **c**, Venn diagram of mouse and human DE pathways affected by neuronal APOE4 — 9 of the top 15 pathways affected in iPSC-derived human excitatory neurons were also found in the 16 DE pathways identified in mouse hippocampal excitatory neurons. These 9 pathways are listed in red. **d**, Gene-pathway network plot of DE genes across the 9 shared DE pathways. Lines link DE genes to each DE pathway of which they are a member. Size of the cluster circle is proportional to number of genes detected in the cluster. Intensity of color of the gene circle reflects upregulation (red) or downregulation (blue) of that gene.

### Human *APOE4* carriers show similarly affected DE genes and pathways in cortical excitatory neurons

We then further compared the transcriptomic effects of APOE4 in mouse excitatory neurons and human iPSC-derived excitatory neurons with those analyzed from a publicly available snRNA-seq dataset (GEO:GSE157827) of human prefrontal cortex from normal control subjects at ages between 74 and 94^39^. Since there was no normal control APOE4/4 carrier available in this cohort, we compared APOE4/3 with APOE3/3 individuals. These human snRNA-seq data were filtered, analyzed, and plotted using Seurat. Marker genes were used to identify the major cell types in the dataset, including excitatory neurons, inhibitory neurons, oligodendrocytes, OPCs, astrocytes, and microglia (Fig. 7a).

**Fig. 7.**
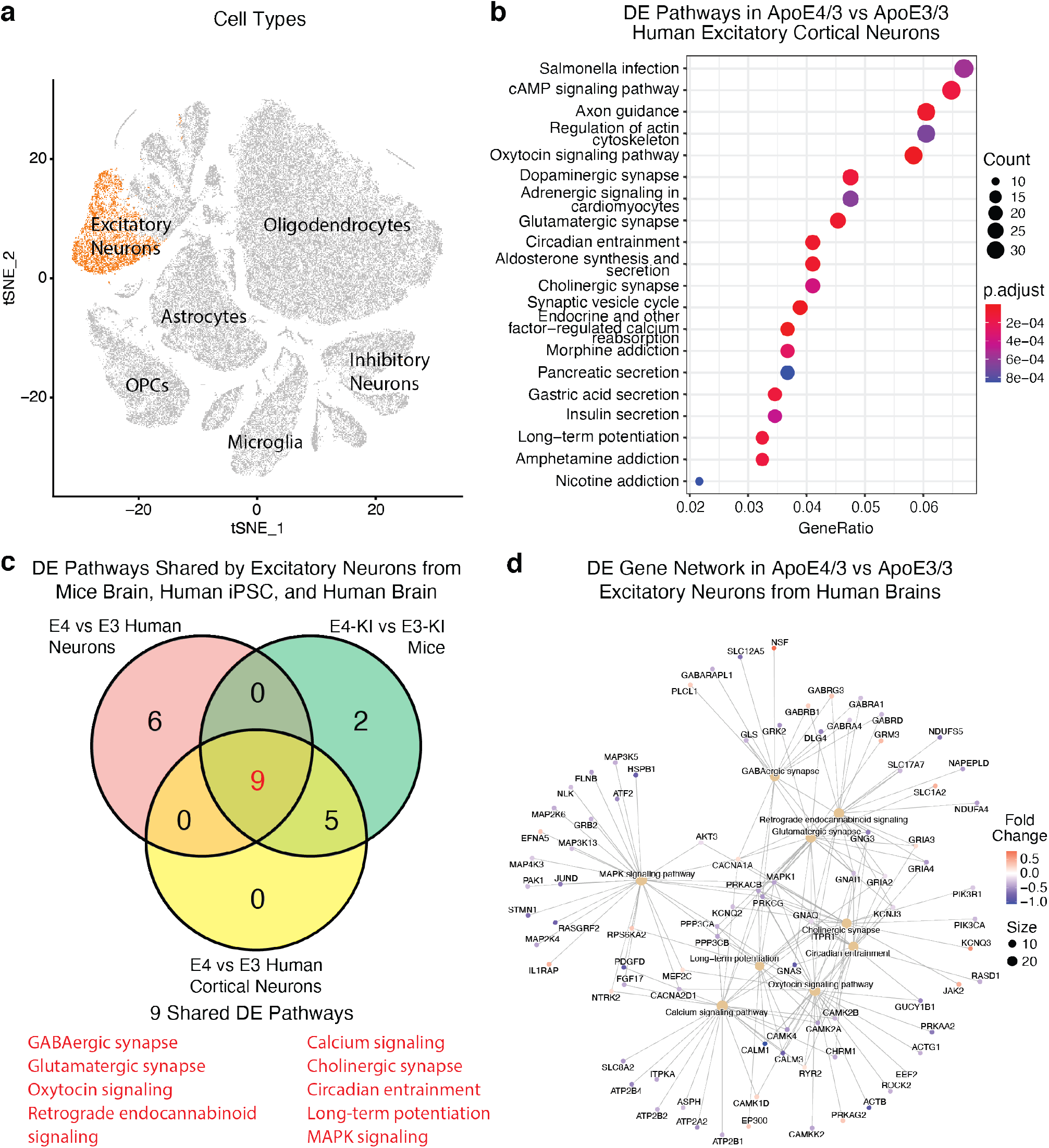
Human *APOE4* carriers show similarly affected DE genes and pathways in cortical excitatory neurons. **a**, tSNE plot shows major cell clusters, including excitatory neurons, inhibitory neurons, oligodendrocytes, OPC, astrocytes, microglia, and others from the analyzed human cortical snRNA-seq dataset (GEO:GSE157827). Excitatory neurons are highlighted in orange and the other major brain cell types are identified by text labels. **b**, Excitatory neurons were analyzed for DE genes between APOE4/3 and APOE3/3 carriers. Top 20 statistically significantly enriched DE pathways are shown. ‘Count’ shows number of DE genes identified in a pathway, corresponding to size of the dot; ‘p.adjust’ shows adjusted P-value for the statistical enrichment of genes in a pathway, red color corresponds to lower P-value. **c**, Venn diagram showing number of DE pathways significantly affected by APOE4 in each combination of mouse excitatory neuron transcriptomic data, human iPSC-derived excitatory neuron transcriptomic data, and human cortical excitatory neuron transcriptomic data. Nine DE pathways were shared by all three transcriptomic datasets. **d**, Gene-pathway network plot of DE genes across the 9 shared DE pathways. Lines link DE genes to each DE pathway of which they are a member. Size of the cluster circle is proportional to number of genes detected in the cluster. Intensity of color of the gene circle reflects upregulation (red) or downregulation (blue) of that gene.

Analysis of the DE genes affected by APOE4 in human cortical excitatory neurons (Supplementary Table 8) identified the top 20 DE pathways (Fig. 7b and Supplementary Table 9). Nine out of the 20 DE pathways in human cortical excitatory neurons overlapped with the 9 DE pathways shared between mouse hippocampal excitatory neurons and human iPSC-derived excitatory neurons (Fig. 7c). Gene network analysis revealed the DE genes across the 9 shared DE pathways in human prefrontal cortical excitatory neurons (Fig. 7d). These data further support consistency in transcriptomic effects of APOE4 among mouse neurons, human iPSC-derived neurons, and human brain neurons.

## Discussion

### APOE4 affects neuronal transcriptomes similarly in mice and humans, starting early and in non-demented individuals

Our initial analyses of APOE4-KI and APOE3-KI mouse hippocampus aimed to identify DE genes and affected pathways in all possible cell types, in mice aged from 5 to 20 months. However, we found the greatest number of APOE4-induced DE genes and pathways in neurons rather than other cell types. Excitatory neurons were the most consistently and strongly affected, beginning at young ages. Therefore, we focused subsequent analyses in this study on APOE4 effects on the transcriptome of excitatory neurons.

The APOE4 effects that we detected in mouse excitatory neurons begin early in life and are prominent at later ages. Similarly, we observed differential gene expression in aged individuals carrying APOE4/3 or APOE3/3 who did not show clinical manifestation of AD. Although these non-demented individuals were aged, the transcriptomic effect of APOE4 in their excitatory neurons suggests that their differences are due to lifelong APOE4 effects rather than resulting from advanced AD. The data of *in vivo* chimeric human neuron model further demonstrate that human excitatory neurons, like mouse excitatory neurons, can show differences based on their own APOE genotype. Our results from mouse excitatory neurons of human APOE-KI mouse lines, human excitatory neurons derived from iPSCs, and human brains show that the effects of APOE4 on excitatory neuronal transcriptome affect common pathways relevant to AD pathogenesis, including synaptic function, calcium signaling, and intracellular signal transduction, across the three data sets. Interestingly, many of these neuronal pathways were also affected by bumetanide, a repurposed drug with potential for preventing or treating APOE4-related AD^40^.

Our snRNA-seq results suggest some causal links between APOE4 expression and known cellular, functional, and behavioral phenotypes. For one prominent example, we found many DE genes and pathways in CA1 neurons of young adult as well as aged mice. CA1 neurons are vulnerable to AD pathologies^41^, neuronal loss^42,43^, dendritic alterations^44^, and loss/alteration of synapses^45^. The APOE4-induced DE genes and affected pathways could contribute to these effects, through early effects on synaptic function, calcium signaling, and intracellular signal transduction — three modules of DE pathways that we identified — as well as responses to disease-related stressors throughout the lifespan.

The expanded number of DE genes detected at later ages, but retention of affected cellular pathways that are already affected at 5 months, suggests that the transcriptomic effects of APOE4 change quantitatively during aging but remain qualitatively somewhat similar. Persistent effects on gene expression throughout life may contribute to cascading changes in neuron development/maintenance, morphology, function, and survival as disease initiates and phenotypes progress.

### Neuronal APOE4 is responsible for the early and persistent effects on neuron transcriptomes

Our snRNA-seq results extend previous studies that have shown impacts of APOE4 expressed in neurons^26,30,31^ as well as astrocytes and other cell types in the brain^31,46^. We detected APOE transcripts in several types of hippocampal neurons, including excitatory and inhibitory neuron clusters, as we reported previously ^31^. Earlier studies have provided other evidence for *APOE* gene expression in neurons, including expression of a GFP-APOE reporter in mouse neurons following injury^29,30^ and transcripts in neuronal cells in vitro using RNAscope, Quantigene, RT-PCR, and RNAseq^47^. The presence of APOE transcripts in neurons suggests the possibility that APOE4 effects on neuronal transcriptomes may be due, at least in part, to APOE4 derived from neurons.

Using the APOE-fKI/Syn1-Cre mice, in which the *APOE* gene is deleted specifically in neurons ^31,37,48^, allows us to test this hypothesis experimentally. We took the strategy of discarding the DE genes if they were still differentially expressed after APOE removal from neurons. This approach shows that, while certain genes are differentially expressed in excitatory neurons regardless of APOE4 expression in neuronal and non-neuronal cells or only non-neuronal cells, most of the DE genes are detected only when APOE4 is present in neuronal cells. Because these DE genes show significant enrichment in the same 16 pathways affected by APOE4 in APOE4-KI versus APOE3-KI mouse hippocampi, this result suggests that the major affected DE genes and pathways are due to endogenous neuronal APOE4 expression. In support of this conclusion, snRNA-seq analysis of the APOE4 and APOE3 human iPSC-derived excitatory neurons isolated after transplantation into APOE4-KI mouse hippocampus^28^ reveals that 9 out of the 16 APOE4-induced DE pathways are shared between the two datasets. Together, these results suggest that APOE4 expression in excitatory neurons is necessary for inducing alterations of the key APOE4-related pathways that we detected. Importantly, the same 9 DE pathways induced by neuronal apoE4 are also identified by comparing the transcriptomes between excitatory neurons carrying APOE4/3 and those carrying APOE3/3 from aged human brain cortex, suggesting the potential for translation of these findings to human brains.

### Key genes and pathways affected by neuronal APOE4

We found consistent neuronal APOE4-induced disruptions across multiple ages and several excitatory neuron types in key KEGG pathways, particularly those related to calcium signaling, synaptic function, and MAPK/Rap1/Pld signal transduction. Importantly, alterations of these pathways have been implicated in AD pathogenesis in studies of animal models and humans.

Disruptions in calcium signaling pathways are found in excitatory neurons from APOE4-KI mouse hippocampus, transplanted human APOE4 iPSC-derived excitatory neurons, and cortical excitatory neurons of human APOE4 carriers. Altered calcium levels inside cells can disrupt neurotransmitter release, postsynaptic signaling, and neuronal activity^49-51^. Our results, in concert with previous data, suggest that calcium signaling disruptions may play an important role in APOE4 effects and start early in the brain. Prior studies have found that APOE4 triggers greater intracellular calcium levels than APOE3, and impacts NMDA-receptor-mediated signaling in cultured neurons to disrupt calcium signaling and protein translation^52^.

Several mechanisms can disrupt calcium levels and calcium signaling in the aging brain^53^ and AD^54^. Calcium interactions with Presenilins and APP may be key mediators in some types of AD^55^. Further research is needed to understand the mechanisms by which APOE4 impacts calcium-regulating gene expression, and the functional consequences of these expression changes.

Alterations of calcium signaling can also interfere with synaptic plasticity and related signaling pathways, contributing to neurodegeneration and memory impairments^56,57^. We found that many genes with critical synaptic functions also had extensive early dysregulation by neuronal APOE4 in the hippocampus. In particular, downregulated genes Grin2a and Grin2b are critical for LTP and LTD, and critical for learning and memory^58,59^. Likewise, Trp6 is very important for dendritic growth and formation of excitatory synapses^60^. Gabra2, GABA_A_ receptor alpha subunit, is involved in generation of brain oscillations critical for cognition^61^. We found a highly significant and consistent downregulation of Gabra2, which could link APOE4 to known oscillatory disruptions^62,63^. Thus, our results support prior studies that have documented synaptic deficits in APOE4-KI mouse brains^32,33,35^ and, importantly, associate these detrimental effects to neuronal APOE4.

Other consistently affected signaling pathways by neuronal APOE4 include MAPK, Rap1, and Phospholipase D. These molecular pathways are functionally related to each other, as well as to calcium and synaptic functions. Both Rap1 (mmu04015) and Phospholipase D (mmu04072) can modulate calcium and MAPK (mmu04010) signaling pathways. Rap1, a small GTPase, can mediate MAPK kinase activation by calcium signaling^64,65^. Rap1 is involved in major aspects of neuron functional morphology, including migration, polarity, axon and dendrite growth, and synapse formation^66,67^. Small GTPases have been linked to AD via regulation of BACE1 and generation of amyloid β^68^. GTPases have also been studied as potential AD drug targets^69^. Although MAPK and Rap1 signaling are closely interconnected, in our analysis the DE genes in mouse MAPK and Rap1 pathways were largely non-overlapping, suggesting that genes in both of these pathways show distinct effects of neuronal APOE4.

### APOE4 affects astrocytic transcriptomes later in life

Consistent with previous studies, we found that APOE expression was primarily localized to astrocytes in mice, with lower levels in neurons and microglia. Our analysis of the human cortex dataset^39^ confirmed that human APOE is expressed at relatively high levels in both astrocytes and microglia compared to neurons. Interestingly, we observed differential gene expression between APOE4-KI and APOE3-KI astrocytes primarily in aged mice (15 months). These differential expression effects of APOE4 in astrocytes may impact neuron health and AD-related phenotypes that are observed by 15 months in APOE4-KI mice. However, they may not be the primary drivers of differential expression effects of APOE4 that emerge earlier in life. It is plausible that the differential gene expression in astrocytes may be affected by the changes that happen over time as neurons experience effects of neuronal APOE4.

APOE4 expression in non-neuronal cell types (astrocytes, microglia, pericytes, etc.) can affect how they interact with neurons^11,12,70^. Bulk RNA-seq analysis revealed that, relative to APOE3, mice with APOE4, which is highly produced in astrocytes, showed significantly more expression of an endopeptidase module including several Serpina genes, and reduced expression of a module related to lysosome and purine deoxyribonucleoside monophosphate metabolism^11^. *APOE*-genotype-dependent effects have been observed in human iPSC-derived astrocytes, where APOE4 astrocytes were less capable of promoting neuronal survival and synaptogenesis^71^. In mice, a human APOE3 transgene was excitoprotective when expressed in either astrocytes or neurons, but APOE4 expression was excitoprotective only in astrocytes. The neuronal APOE4 transgene was not excitoprotective and contributed to neuronal loss following excitotoxic injury with kainic acid^72^. Deletion of APOE4 specifically in neurons using the APOE-fKI/Syn1-Cre line protected mice from GABAergic interneuron loss and memory deficits, while deletion of APOE4 in astrocytes did not show neuronal and behavioral protections^48^. Thus, neuronal APOE4 appears to possess gain-of-toxic-function effects directly on neurons, which occur early in life. Astrocytic APOE4 seems to have loss-of-function effects on astrocytes’ neuronal protective roles, which occurs later in life. These possibilities warrant further studies.

Our large-scale snRNA-seq study with human APOE-KI mice identified prominent effects of neuronal APOE4 on neuronal transcriptomes related to synaptic function, calcium signaling, and intracellular signal transduction, already present by 5 months of age and persisting during aging. Importantly, APOE4 affects the expression of similar genes and pathways in human APOE4 iPSC-derived neurons transplanted into APOE4-KI mouse hippocampus and in human neurons from aged brains, supporting the potential of translating these findings to humans. Further work is needed to understand the contribution of these gene expression changes to disease initiation and/or progression in specific cell types. Furthermore, these data provide a foundation for developing drugs targeting neuronal APOE4 and indicate the requirement of early interventions for successfully treating APOE4-related AD.

## Materials and Methods

### Mice

All protocols and procedures followed the guidelines of the Laboratory Animal Resource Center at the University of California, San Francisco (UCSF) and the ethical approval of the UCSF institutional animal care and use committee (IACUC). Experimental and control mice had identical housing conditions from birth through sacrifice (12 h light/dark cycle, housed 5 animals/cage, PicoLab Rodent Diet 20). APOE3-KI and APOE4-KI homozygous mouse lines (Taconic)^73^ were born and aged under normal conditions at the Gladstone Institutes/UCSF animal facility. APOE4-fKI/Syn1-Cre and APOE3-fKI/Syn1-Cre mice were generated by cross-breeding APOE-floxed-KI mice, which was generated in our lab^37^, with Syn1-Cre mice^48^.

### cDNA library preparation

cDNA libraries were prepared using the 10x Chromium Single Cell 3’ GEM, Library & Gel Bead Kit according to the manufacturer’s instructions. 10x Chromium v2 kit was used for the APOE-KI dataset. 10x Chromium v3 kit was used for APOE-fKI/Syn1-Cre dataset (10x Genomics: 1000092). Libraries were sequenced on an Illumina NovaSeq 6000 sequencer at the UCSF CAT Core.

### Pre-processing and clustering of mouse snRNA-seq samples

For the APOE-KI mouse data, samples were processed as previously described^31^. Briefly, demultiplexed FASTQ files were aligned to a custom reference genome built from mm10-1.2.0 that includes introns, using the Cell Ranger v2.0.1 counts function with default parameters, as described in the Cell Ranger documentation. For APOE-fKI/Syn1-Cre mice, introns were included by using mm10 genome with cellranger-4.0.0 and the include-introns option.

UMI counts were also determined using the Cell Ranger counts function, and count matrices from different samples were aggregated into a single count matrix using Cell Ranger’s aggr function with default parameters. This function filters barcodes (potential cells) according to a UMI count threshold. Filtered UMI count matrices were further processed using Seurat. Cells were filtered to include only cells with 200–2,400 genes detected, 500–4,500 UMIs and <0.25% mitochondrial reads. This quality assurance process resulted in a final matrix of 21,204 genes by 123,489 nuclei. The gene expression matrices were then log-normalized with a scale factor of 10,000, using the Seurat NormalizeData function^74,75^.

Highly variable genes were calculated via the FindVariableGenes function with default parameters. The expression level of highly variable genes in the nuclei were scaled and centered via the ScaleData function, and was then fed into the RunPCA function.

### Cell clustering

For the APOE-KI mouse dataset, existing clustering from a published study was used ^31^. For APOE-fKI/Syn1-Cre mice, transplanted neurons derived from human iPSCs, and human cortical neuron data, we filtered and clustered snRNAseq data in R. Data visualization by tSNE revealed clusters where mouse ages and genotypes were intermingled, with no discernable evidence of batch effects by genotype or age.

Seurat embeds cells in a *k*-nearest neighbor graph, based on Euclidean distance in PCA space. The edge weights between any two cells are further refined using Jaccard similarity. Highly dispersed genes were selected using the Seurat FindVariableGenes function^57,58^. The expression level of highly variable genes in the nuclei were scaled and centered via the ScaleData function, and was then fed into the RunPCA function. Nearest neighbor distances were computed using up to the first 15 PCs and a resolution of 0.6.

### Cell type assignment

Marker genes for each cluster were calculated using the FindAllMarkers function in Seurat^74,75^. This algorithm iteratively compares gene expression in each putative cluster against the expression in all other clusters, using the Wilcoxon rank-sum test. Marker genes were limited to be positively expressed (more highly expressed in the cluster of interest than in other clusters), to be detected in at least 10% of cells in the cluster and to be 0.25 log2 fold higher expressed in the cluster of interest than in other clusters.

Broad cell classes, such as excitatory and inhibitory neurons, astrocytes, oligodendrocytes and OPCs, were identified by querying marker genes against cell-type-specific markers derived from previous RNA sequencing data on sorted cell types, publicly available online at brainrnaseq.org^76^. For further subdivision of hippocampal cell types, particularly for identification of subsets of principal cells, marker genes for each cluster were queried against hippocampal cell-type-specific marker genes as published from hipposeq.org^77^.

For comparison of APOE-KI with APOE-fKI/Syn1-Cre DE genes, we compared the DE gene data from 15-month-old mice APOE-KI (clusters 1, 2, 3, 6, 10, 12, 17, 19, 21, 26, 27) with those from 15-month-old APOE-fKI/Syn1-Cre mice (clusters 1, 2, 3, 6, 10, 11, 14, 15, 17, 21, 23) ^31^. The 5-month-old and 10-month-old mice comparisons were done separately, for which we compared APOE-KI DE genes from cell clusters 1, 2, 3, 6, 9, 10, 17, 19, 21, 24, 25, 26, 27 with APOE-fKI/Syn1-Cre DE genes from cell clusters 1, 2, 3, 11, 6, 8, 13, 16, 18, 20, 24, 26, 28 from a new dataset.

### hiPSC-derived neuron data

Generation of hiPSC-derived neurons, transplantation, single nuclei FACS sorting of transplanted human neurons from mouse hippocampi, human cDNA library preparation and sequencing, and pre-processing and clustering of human snRNA-seq samples were as described in detail previously^28^.

### Human subjects

Anonymized human cortical snRNA-seq data were downloaded from GEO:GSE157827 of human prefrontal cortex from normal control subjects at ages between 74 and 94, with APOE genotypes confirmed by TaqMan assay^39^. These data included the output files from Cell Ranger (version 3.0.1) with the default settings (barcodes.tsv.gz, matrix.mtx.gz, and features.tsv.gz). We loaded this data into Seurat for clustering. Cells were filtered to include only cells with 200–2,400 genes detected, 500–4,500 UMIs and <2.5% mitochondrial reads. Clinical and demographic data from the human snRNA-seq cohort can be found in Supplementary Table 1 of ref ^39^.

### Cell counts

Two-way ANOVA tests were conducted with the aov() function in R. First, for each genotype at each age, the percentage of nuclei in each cluster was calculated. The ANOVA was then used to test for effects of age, genotype, and age:genotype interaction (*y ∼ APOE genotype × age*) on this percentage of nuclei for each cluster.

### Differential gene expression analysis

We used the zinbwave R package to process the snRNA-seq data from each cluster^78^ and edgeR to test differential expression^79^. The zinbwave package calculates observational weights to obtain a low-dimensional representation of the data. Each cluster was analyzed separately. Genes were removed from the analysis if they did not have more than 1 reads in greater than 1% of samples. The zinbwave function was used with specified parameters K=2, epsilon=1000, observationalWeights = TRUE. Differential expression was calculated with the edgeR package using estimateDisp(), glmFit(), and glmWeightedF() functions. The Benjamini-Hochberg method was used to control the false discovery rate.

### KEGG pathway enrichment analysis

KEGG (Kyoto Encyclopedia of Genes and Genomes) is a collection of manually drawn pathway maps representing molecular interaction and reaction networks. Enriched KEGG pathways for heatmaps were predicted using clusterProfiler package in R^80^, which uses data from the online KEGG mouse pathway database. The over representation analysis uses a hypergeometric distribution to calculate whether KEGG pathways are overrepresented in a list of differentially expressed genes. For KEGG Gene-Pathway Heatmaps, genes were included as differentially expressed genes if false discovery rate (FDR) < 0.05 and logFC > 0.1. We used the enrichKEGG() function, and included KEGG pathways with minGSSize (minimum number of genes in a gene set) = 10, pvalueCutoff (adjusted p-value cutoff) = 0.05, and was filtered using gsfilter() from the DOSE package with ‘Count’ min (minimum number of differentially expressed genes present in a gene set) = 4. FDR for enriched pathways was calculated using Benjamini & Hochberg correction. Gene-pathway heatmap graphs were generated using the heatmap.2() gplots function, with colors representing logFC and dendrogram clustering for both genes and KEGG pathways. Pathway network visualizations were conducted in Cytoscape^81^.

## Data Availability

The APOE3-KI and APOE4-KI mouse snRNA-seq data used in this study are available from the Gene Expression Omnibus (www.ncbi.nlm.nih.gov/geo) with an accession number GSE167497. These data are used for Figures 1, 2, and 3, as well as for comparisons in subsequent figures.

Additional sequencing data of APOE3-fKI/Syn1-Cre and APOE4-fKI/Syn1-Cre mice, generated in this study, will be submitted to Gene Expression Omnibus and made publicly available.

Data from snRNA-seq of in vivo chimeric mice are available in the Gene Expression Omnibus (accession number: GSE152867)

Single-cell RNA-seq data from human brain, referenced in Figure 7, are available in the Gene Expression Omnibus (accession number: GSE157827).

The Kyoto Encyclopedia of Genes and Genomes Pathways (KEGG) database is available at https://www.genome.jp/kegg/pathway.html.

## Code Availability

The custom scripts used for analyzing data and creating plots during this study will be made publicly available on Github during the review process.

## Acknowledgements

This work was partially supported by grants RF1AG055421, R01AG071697, P01AG073082 to Y.Huang from the National Institutes of Health (NIH). The results published here are, in part, based on data obtained from GEO:GSE167497, GEO:GSE152867, and GEO:GSE157827. We thank Alice Taubes for providing helpful discussions about snRNA-seq and data analysis in R. The Gladstone Flow Cytometry Core FACSAria cell sorter is supported by NIH S10 RR028962 and the James B. Pendleton Charitable Trust. We thank Eric Chow and the staff at the UCSF CAT Core for advice and support with RNA sequencing; Nandhini Raman of the Gladstone Flow Cytometry Core for sorting the nuclei; Steve Belunek and Wil Maguire of Gladstone Information Technology for server support; Reuben Thomas at Gladstone Bioinformatics Core for assistance with snRNA-seq analysis; and Theodora Pak for editorial assistance.

## Author contributions

B.P.G. and Y.Huang designed and coordinated the study and wrote the manuscript. B.P.G. carried out most studies and data analysis. K.A.Z. generated the original snRNA-seq dataset (GEO:GSE167497) from APOE-KI mouse hippocampi at different ages and helped on snRNA-seq data analysis. Y.Hao dissected mouse hippocampi, isolated cell nuclei, and prepared samples for RNA sequencing. S.Y.Y. and P.A. managed mouse lines and brain collections. Y.Huang supervised the project.

---

**Extended Data Fig. 1.**
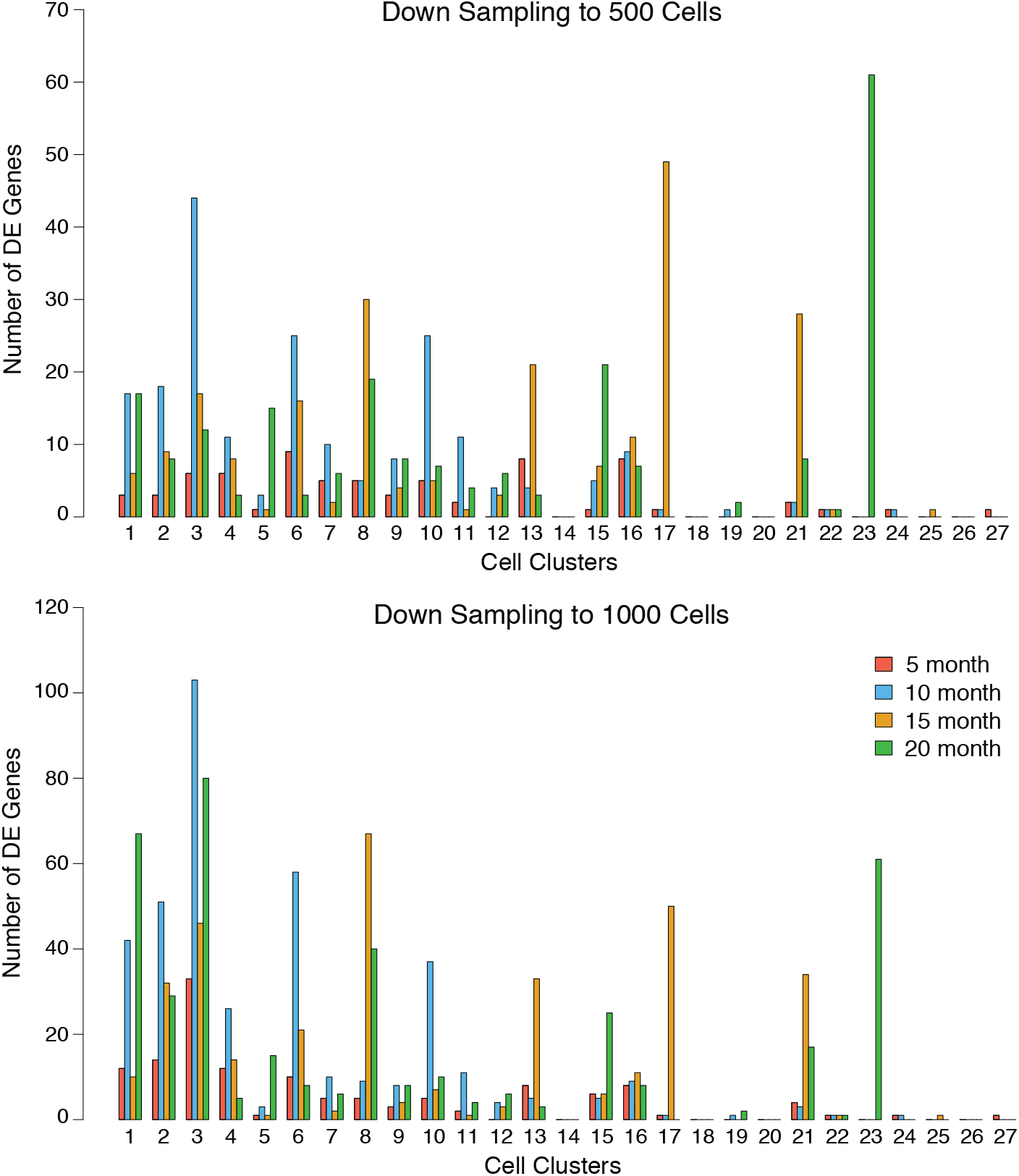
Down-sampling the APOE-KI mouse data at each age shows consistent effects of genotype, regardless of nuclei sampling size. **a**,**b**, The number of DE genes between APOE4-KI and APOE3-KI samples is shown on the Y-axis for down-sampling to 500 cells (a) or 1,000 cells (b). The X-axis shows data separately for each cell cluster.

**Extended Data Fig. 2.**
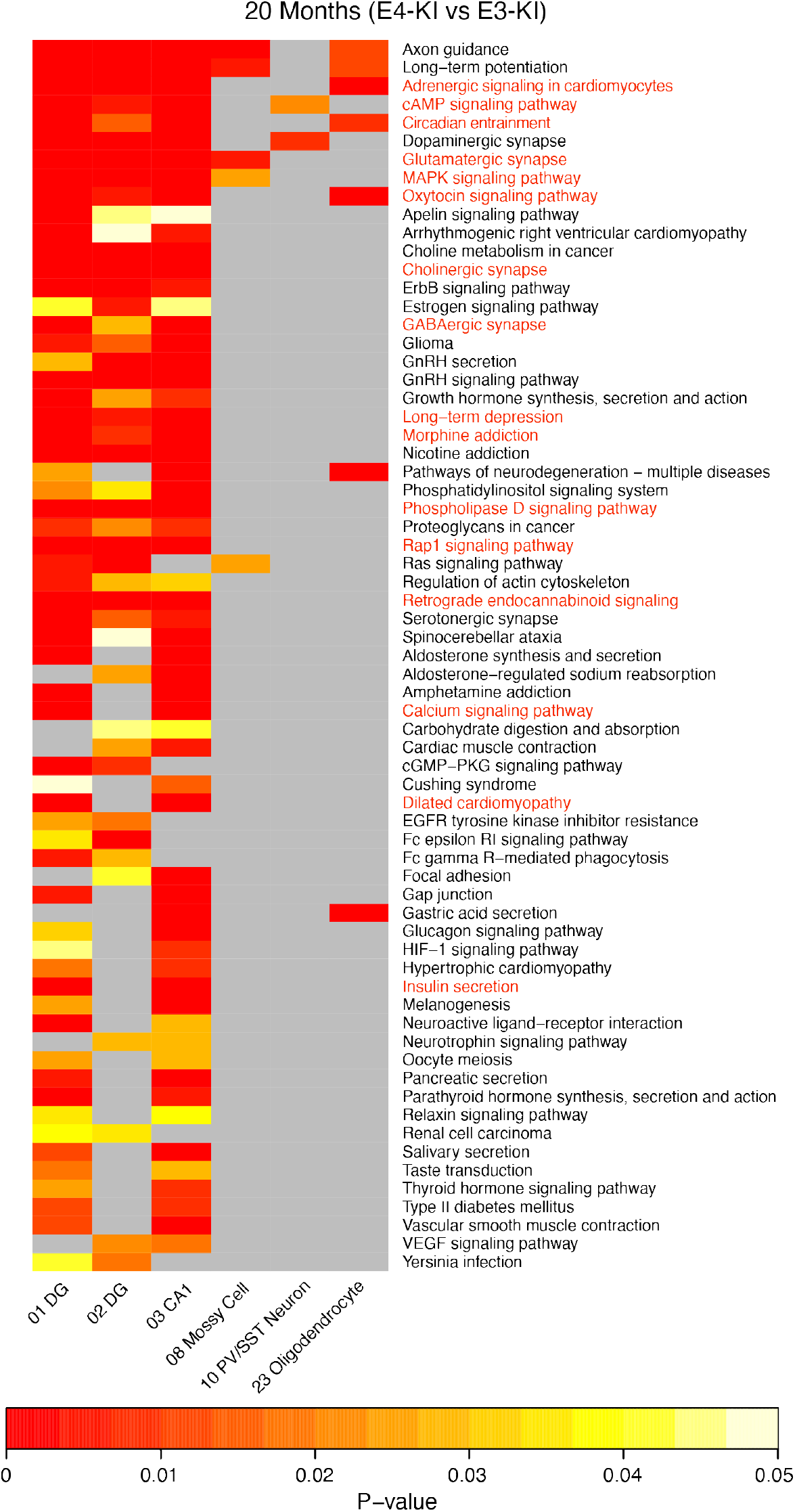
APOE4 induced DE pathways in hippocampal neurons at 20 months of ages. DE genes of APOE4-KI versus APOE3-KI mouse hippocampal cell types are significantly enriched in several KEGG pathways at 20 months of age. Heatmap plots show cell clusters with significantly enriched DE pathways (on the x-axis) and the top enriched DE pathways (on the y-axis) sorted from most commonly detected (top) to least (bottom). P-value for statistical enrichment is shown by color, with red indicating smaller P-value, gray indicating P-value > 0.05, and scale indicated in the legend. Pathway names highlighted in red are found at all ages.

**Extended Data Fig. 3.**
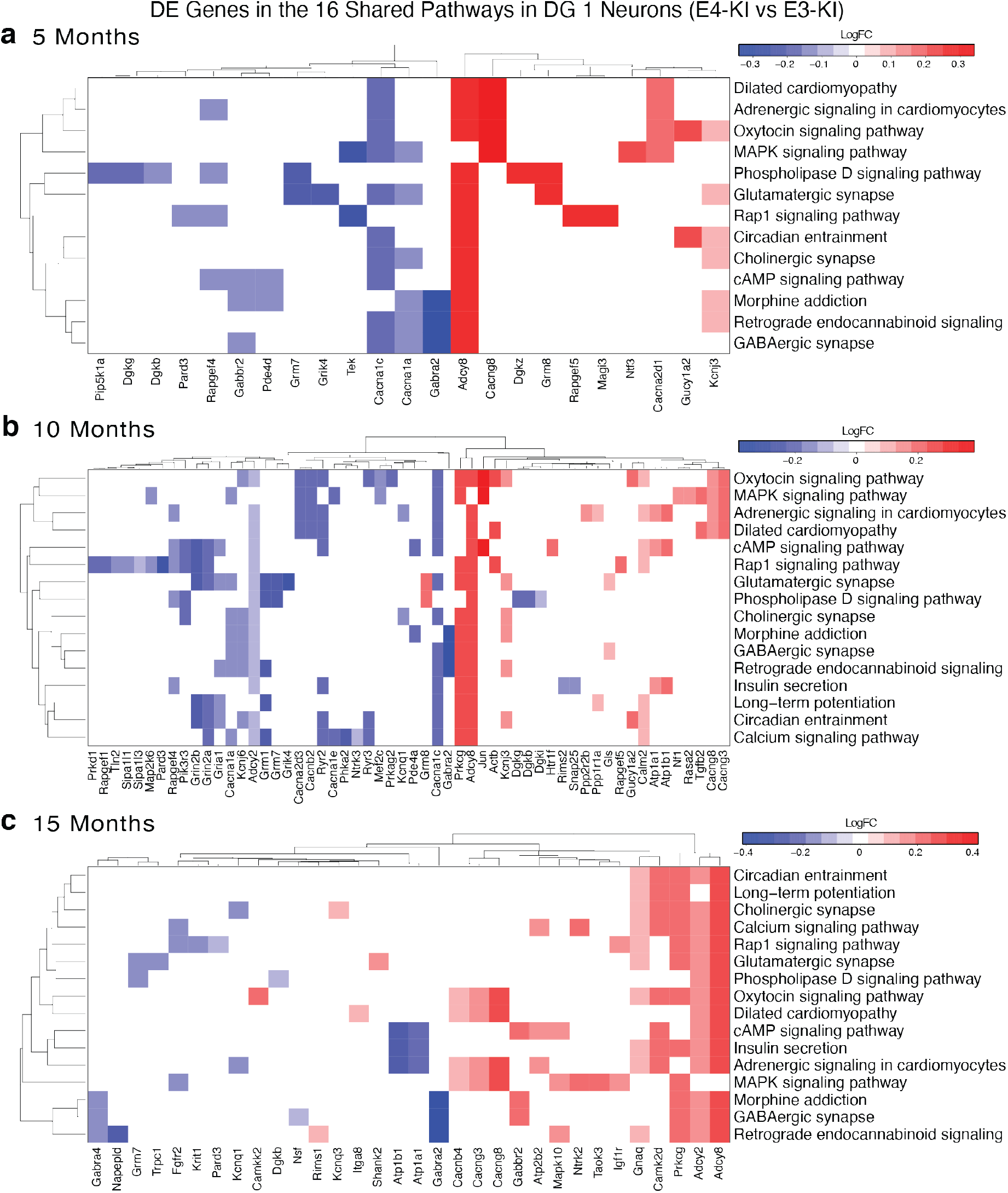
APOE4-induced DE genes in the 16 shared DE pathways in hippocampal dentate gyrus granule cells (cluster 1) across age. **a–c**, The DE genes in 16 shared pathways affected by APOE4 were plotted as heatmaps at 5 months (a), 10 months (b), and 15 months (c). For this analysis, we used the enrichKEGG() function, and included KEGG pathways with minGSSize = 10, pvalueCutoff = 0.05, Count min = 4. Genes were included if FDR < 0.05 and logFC > 0.1. The APOE4 effect, relative to APOE3, is plotted as intensity of blue (downregulation) or red (upregulation), scale indicated in the legend. DE genes are clustered on the x-axis,and pathways are clustered on the y-axis, by similarity of expression effects.

**Extended Data Fig. 4.**
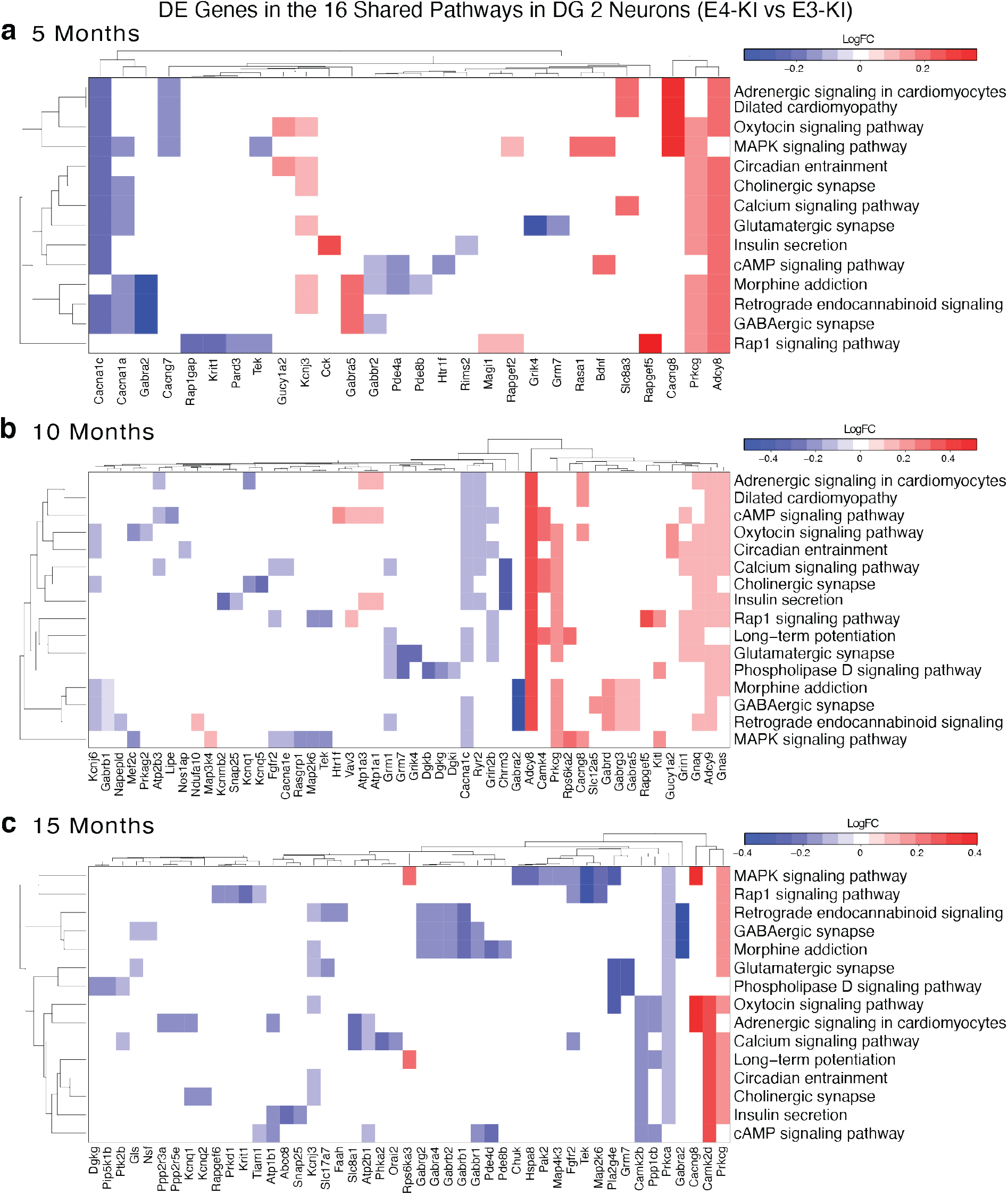
APOE4-induced DE genes in the 16 shared DE pathways in hippocampal dentate gyrus granule cells (cluster 2) across age. **a–c**, The DE genes in 16 shared pathways affected by APOE4 were plotted as heatmaps at 5 months (a), 10 months (b), and 15 months (c). For this analysis, we used the enrichKEGG() function, and included KEGG pathways with minGSSize = 10, pvalueCutoff = 0.05, Count min = 4. Genes were included if FDR < 0.05 and logFC > 0.1. The APOE4 effect, relative to APOE3, is plotted as intensity of blue (downregulation) or red (upregulation), scale indicated in the legend. DE genes are clustered on the x-axis,and pathways are clustered on the y-axis, by similarity of expression effects.

**Extended Data Fig. 5.**
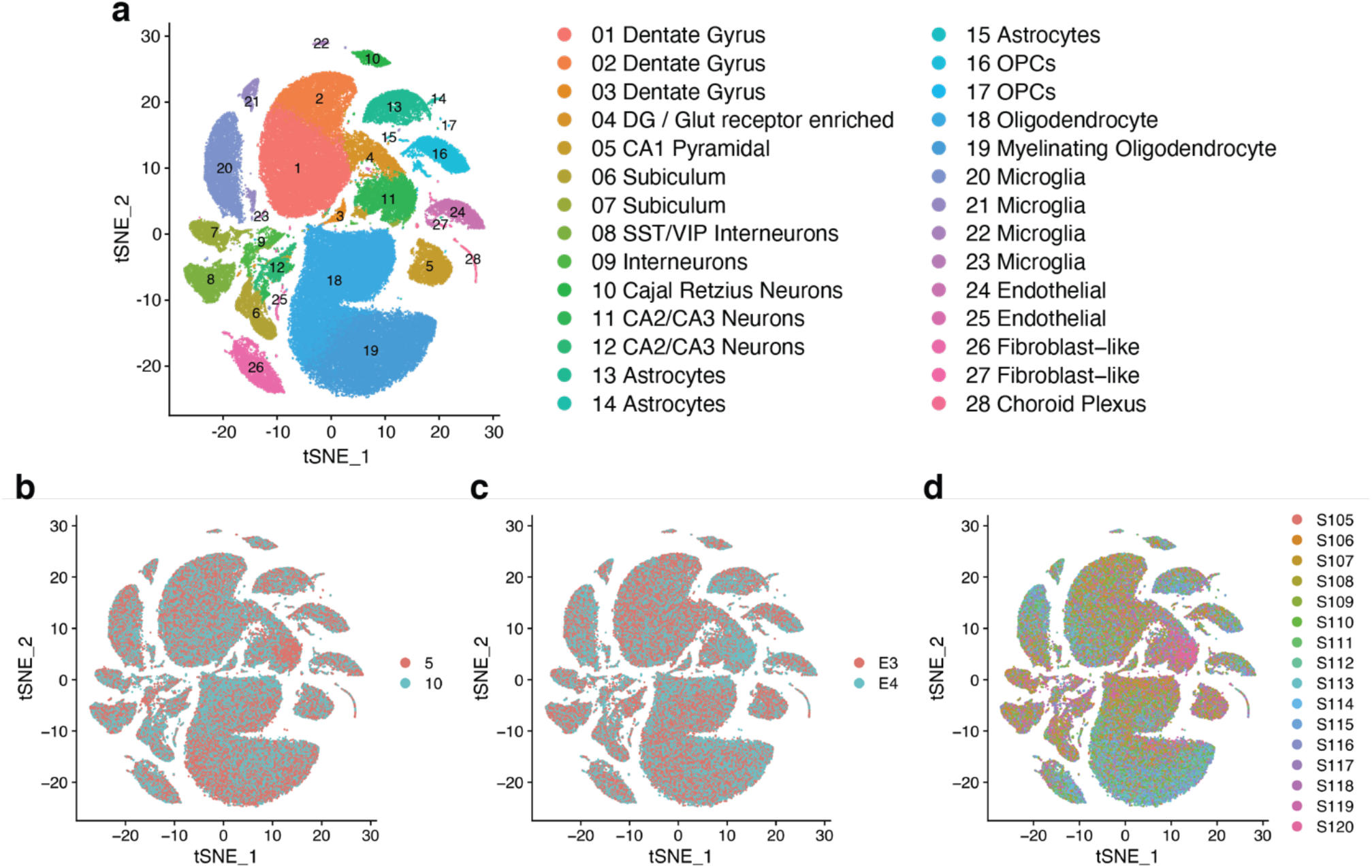
Generation of an APOE-fKI/Syn1-Cre data set for analysis of neuron-specific APOE effects. Hippocampal snRNA-seq data were collected from APOE4-fKI/Syn1-Cre and APOE3-fKI/Syn1-Cre mice at 5 and 10 months (n = 4 mice per genotype per age, total 16 mice). The hippocampi were dissociated, and nuclei were labeled with DAPI and isolated using flow cytometry before processing using the 10x Chromium v3 system. **a**, The snRNA-seq data were filtered, normalized, and clustered using the Seurat package, which yielded 28 distinct cellular clusters, including 12 neuronal clusters (Clusters 1–12) and 16 non-neuronal clusters (Clusters 13–28). Cell type identities for each cluster were identified using positively enriched marker genes. The tSNE plot shows numbered clusters; the corresponding key shows cell types for each numbered cluster. **b–d**, Single nuclei are distributed across all 28 clusters in samples from both ages (b), APOE genotypes (c), and all 16 individual mice (d).

**Extended Data Fig. 6.**
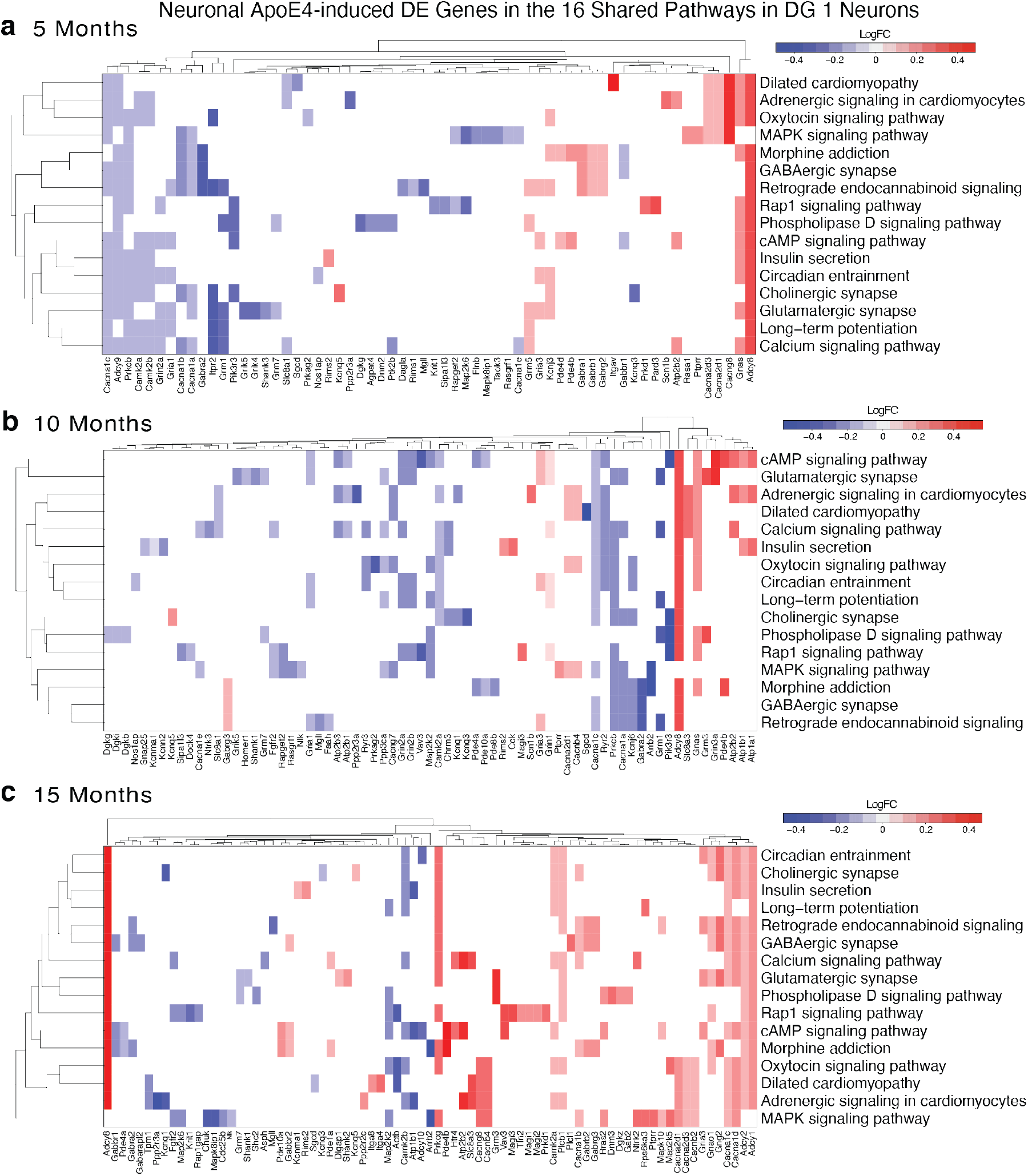
Neuronal APOE4-induced DE genes in hippocampal dentate gyrus granule neurons across age. **a–c**, The cluster 1 neurons (DG1) revealed genes that were differentially expressed in APOE4-KI versus APOE3-KI mice, but not in APOE4-fKI/Syn1-Cre versus APOE3-fKI/Syn1-Cre mice. These neuronal APOE-affected DE genes that fell into 16 consistently affected pathways were plotted as heatmaps at 5 months (a), 10 months (b), and 15 months (c). For this analysis, we used the enrichKEGG() function, and included KEGG pathways with minGSSize = 10, pvalueCutoff = 0.05, Count min = 4. Genes included were DE genes between APOE4-KI and APOE3-KI mice, with genes removed if they were also DE genes between APOE4-fKI/Syn1-Cre and APOE3-fKI/Syn1-Cre mice. For each DE gene, the expression effect of neuronal APOE4 versus APOE3 was plotted as intensity of blue (downregulation) or red (upregulation).

**Extended Data Fig. 7.**
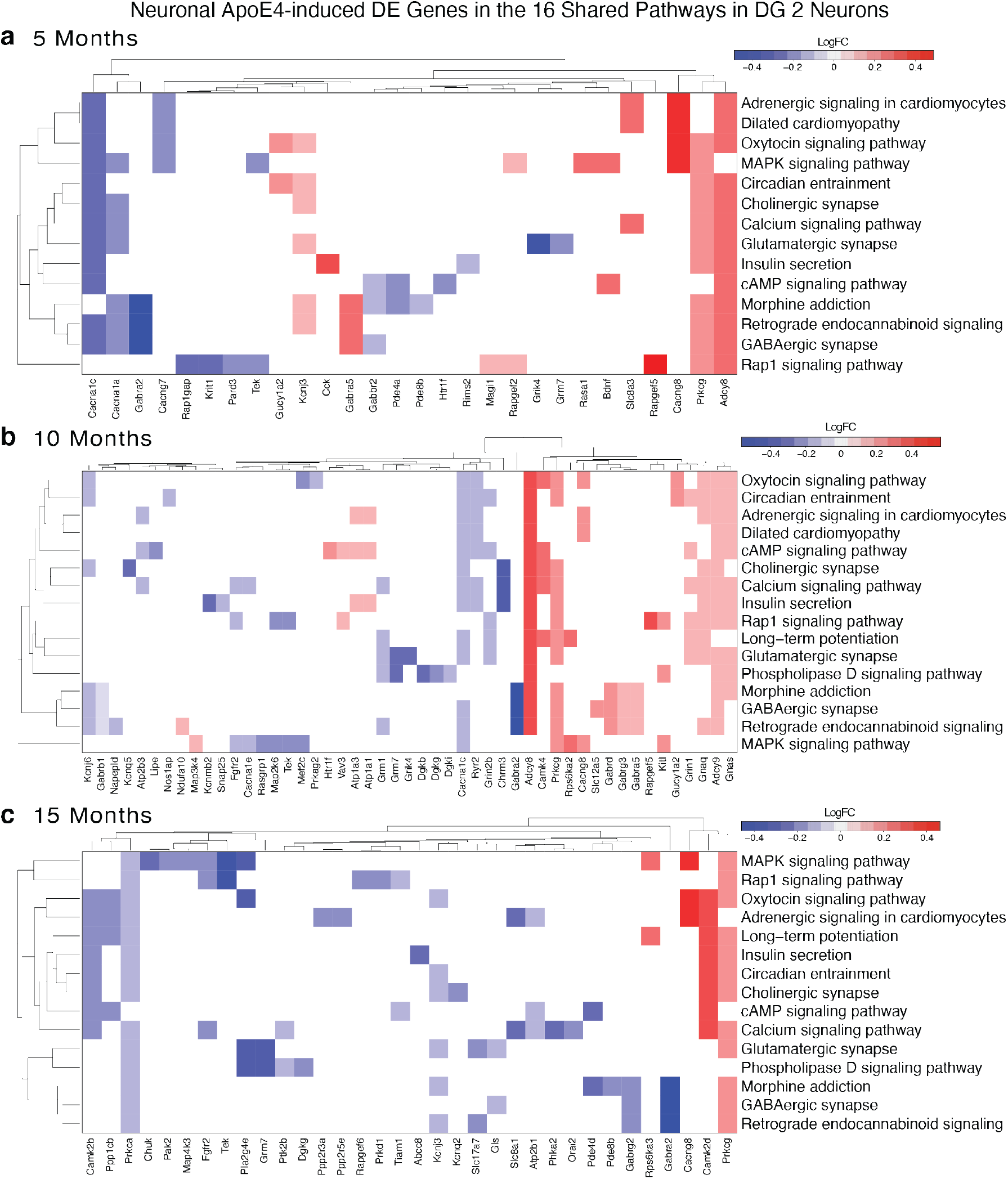
Neuronal APOE4-induced DE genes in hippocampal dentate gyrus granule neurons (DG2) across age. **a–c**, The cluster 2 neurons (DG2) revealed genes that were differentially expressed in APOE4-KI versus APOE3-KI mice, but not in APOE4-fKI/Syn1-Cre versus APOE3-fKI/Syn1-Cre mice. These neuronal APOE-affected DE genes that fell into 16 consistently affected pathways were plotted as heatmaps at 5 months (a), 10 months (b), and 15 months (c). For this analysis, we used the enrichKEGG() function, and included KEGG pathways with minGSSize = 10, pvalueCutoff = 0.05, Count min = 4. Genes included were DE genes between APOE4-KI and APOE3-KI mice, with genes removed if they were also DE genes between APOE4-fKI/Syn1-Cre and APOE3-fKI/Syn1-Cre mice. For each DE gene, the expression effect of neuronal APOE4 versus APOE3 was plotted as intensity of blue (downregulation) or red (upregulation).

